# ABSCISIC ACID INSENSITIVE 5 fine-tunes expression of lipid body genes through modulating ABSCISIC ACID INSENSITIVE 3 binding strength

**DOI:** 10.64898/2026.09.12.751185

**Authors:** Cheuk Him Jack Chiang, Milad Alizadeh, Liang Song

## Abstract

Seed development and germination are two important transitions in a plant’s life cycle. During such transitions, extensive transcriptional regulation reprogramming is performed to support major changes in cellular, tissue, and physiological levels. Transcription factors contain diverse interacting partners for precise transcriptional regulations. Different combinations of interactors would affect temporal and spatial transcriptional regulations. While a large number of transcription factors (TFs) are involved in these transcriptional networks, *in vivo* investigation on how TF interaction dynamics affect transcriptional regulation in seed-to-seedling transition is lacking. ABSCISIC ACID INSENSITIVE 3 (ABI3) has played a major role serving as the center hub of these transcriptional networks during seed development. ABI3 has diverse interacting partners, including ABI5. The presence of these interacting partners allows ABI3 to bind to diverse cis-regulatory elements (CRE) in a precise manner. One of the processes collectively governed by ABI3 and ABI5 is nutrient deposition, including lipid storage. ABI3 and ABI5 regulate lipid body (LB) related genes such as oleosins for controlling LB dynamics. Notwithstanding, not much is known on ABI5’s significance on ABI3’s binding towards CREs. Here, we showed that ABI5 impacts ABI3’s binding strength towards G-Box motifs. The interaction of ABI3 and ABI5 strengthens ABI3’s binding towards ABI3 induced genes. We further demonstrated that ABI3 and ABI5 regulates far more LB related genes during seed development and germination, suggesting that ABI3 and ABI5’s have a more precise role in regulating LB dynamics. Our results had further consolidated ABI5’s role in ABI3’s binding preferences and strength and its significance in controlling LB during seed development and germination.

## Introduction

Seed plants evolved more than 350 million years ago (Linkies et al., 2010). As seed plants evolved, reproductive propagules shifted from predominantly unicellular spore structures to complex multicellular seed structures that provides intricate mechanisms for nutrient deposition and protection (Baroux & Grossniklaus, 2019; Linkies et al., 2010). ABSCISIC ACID (ABA) is one of the hormones that are involved, through diversified mechanisms, in propagule development throughout land plants (Kua et al., 2026). In mosses, ABA promotes formation of brachycytes through reprogramming transcriptomic and cellular activity, facilitating assexual reproduction (Arif et al., 2019; Jinno et al., 2025). In pteridophytes, ABA is involved in sex determination through controlling meristem speciation and division, a crucial step for sexual reproduction (McAdam et al., 2016; Yang et al., 2025). In spermatophytes, ABA promotes seed development including embryogenesis, acquisition of desiccation tolerance and dormancy, and nutrient deposition (Gazzarrini & Song, 2024). While ABA’s role in propagule development diverges, it contains conserved roles in acquiring desiccation tolerance and dormancy, ensuring survival of land plants under stresses (Cuming & Stevenson, 2015; Sussmilch et al., 2017). Seeds contained additional nutrients compared to spores, in the forms of starch, lipid, or proteins (Baroux & Grossniklaus, 2019). While ABA contributes to nutrient deposition during seed maturation, its involvement in reserving nutrients in spores are not fully understood.

The canonical ABA signalling pathway initiates when ABA binds to ABA receptors PYRABACTIN RESISTANCE/PYRABACTIN RESISTANCE 1-LIKE/REGULATORY COMPONENT OF ABA RECEPTOR (PYR/PYL/RCAR) (Ng et al., 2014; Nishimura et al., 2009; Yin et al., 2009). The binding recruits PROTEIN PHOSPHATASE TYPE 2C (PP2C), derepressing SUCROSE NON-FERMENTING-1-RELATED KINASE 2 (SnRK2) for phosphorylation (Ma et al., 2009; Park et al., 2009; Umezawa et al., 2009). SnRK2s thus further phosphorylates downstream transcription factors (TFs) for transcriptional activation (Fujii & Zhu, 2009; Fujita et al., 2009; Nakashima et al., 2009). ABSCISIC ACID INSENSITIVE 3 (ABI3) is one of the TFs downstream of the ABA signalling pathway which is conserved in land plants. ABI3’s physiological significance in dormancy and desiccation tolerance is also conserved throughout land plants. For example, ABI3 orthologs in *Marchantia polymorpha* regulate gemmae dormancy (Eklund et al., 2018), whereas ABI3 in higher plants participates in dormancy acquisition during seed development (Nambara et al., 1992). ABI3 also regulates LATE EMBRYOGENESIS ABUNDANT (LEA) proteins in mosses, Marchantia, legumes, maize, and Arabidopsis (Bedi et al., 2016; Delahaie et al., 2013; Eklund et al., 2018; Khandelwal et al., 2010; Sakata et al., 2010; Thomann et al., 1992). ABI3 is one of the main TFs orchestrating late maturation processes in seeds, including chlorophyll degradation, acquiring desiccation tolerance and dormancy, and nutrient deposition in the form of lipids and proteins (Gazzarrini & Song, 2024). Mutations in ABI3 proteins disrupt these processes during the late maturation stage, leading to green embryos, desiccation intolerant, nondormant, and disrupted nutrient reservoir in spermatophytes (Delmas et al., 2013; Gazzarrini & Song, 2024; Suzuki et al., 2001; Zeng & Kermode, 2004).

Developmental transitions require complex transcriptional regulation. Eukaryotic TFs interact with diverse interacting partners for fine tuning transcriptional activity during such transitions. One classic example is OCTAMER-BINDING TRANSCRIPTION FACTOR 4 (OCT4) and SEX-DETERMINING REGION Y-BOX TRANSCRIPTION FACTOR 2 (SOX2) in mammalian pluripotent stem cells (Mistri et al., 2015; Xie et al., 2025). The presence of SOX2 alters DNA structures that promote OCT4 binding (Mistri et al., 2015; Xie et al., 2025).

Interactions between SOX2 and OCT4 promotes OCT4 binding stability towards DNA regions (Mistri et al., 2015; Xie et al., 2025). Similar mechanisms have also been demonstrated in plant systems. For instance, LEAFY COTYLEDON 1 (LEC1) exhibits combinatorial interactions with diverse TFs for regulating different gene expressions throughout seed development in soybean by binding to different DNA motifs (Jo et al., 2020). ABI3, a B3 domain-containing TF that binds to the RY/Sph element (CATGCA) (Tian et al., 2020), contains diverse interacting partners that allows fine tuning of gene expression levels at different temporal and spatial occasions.

Through binding with different groups such as basic Leucine Zipper (bZIP) TFs, ABI3 gains the ability to differentially bind and activate different genes. For example, ABI3 and ABI5 binds to promoter EM-LIKE1 (EM1) for expressing LEA proteins (Sakata et al., 2010); bZIP10/25 heterodimer interacts with bZIP53 and ABI3 for activation of storage protein genes as nitrogen and sulphur reservoirs (Alonso et al., 2009). On the other hand, ABI3 and ABI5 collectively participate in deposition of lipid storage through regulating DIACYLGLYCEROL ACYLTRANSFERASES (DGATs) for production of neutral lipids like triacylglycerols (TAGs) and oleosins for accumulation of TAGs and stabilisation of lipid bodies (LBs) (Chen et al., 2014; Crowe et al., 2000; Kong et al., 2013; Lee et al., 2022; McGuire et al., 2025). They collectively regulate the formation of LB during seed development, providing an energy source in the future germination stages.

Lipid bodies, also known as lipid droplets, oil bodies and oleosomes, are mono-layered membrane organelles that are known to store nutrients in the form of lipids such as TAGs and sterol esters during stress (Guzha et al., 2023). LBs are initiated in the endoplasmic reticulum (ER), where TAGs and sterol esters are synthesized and accumulated between ER membrane leaflets (Guzha et al., 2023). At the same time, structural proteins such as oleosins and caleosins are co-translationally inserted at LB formation sites in the ER (Guzha et al., 2023). As LB precursors mature, ER resident protein SEIPIN1, together with other LB-associated coating and adaptor proteins facilitates the budding of LB (Guzha et al., 2023). LB dynamics and metabolism are carefully governed by numerous coating proteins, including oleosins, caleosins, OIL BODY-ASSOCIATED PROTEINS (OBAPs), LIPID DROPLET PROTEIN OF SEEDS (LDPS), and SUGAR-DEPENDENT 1 (SDP1) (Guzha et al., 2023). While oleosins and OBAPs are responsible for prevention of LB fusion (Lopez-Ribera et al., 2014; Shimada & Hara-Nishimura, 2010; Siloto et al., 2006), LDPS are known to promote LB fusion during early stages of seed germination (Doner et al., 2025). SDP1, on the other hand, is known to promote degradation of storage lipids via degrading TAGs and trafficking fatty acid (FA) to glyoxysomes (Kelly et al., 2011; Thazar-Poulot et al., 2015)

Although it is known that ABI3 and ABI5 regulated LBs through controlling gene expressions of oleosins and caleosins during seed development and germination, how ABI3 and ABI5 orchestrates LB dynamics in a precise manner is not known. Here, we created *abi5* mutants in ABI3 attached with epitope tags and performed chromatin-immunoprecipitation coupled with sequencing (ChIP-seq) to investigate if ABI3 exhibited differential binding patterns under the influence of ABI5. We further evaluated proteins related to LB dynamics on whether they are regulated by ABI3 and ABI5 and studied if LB dynamics are affected by ABI3 and ABI5 through microscopy. Through performing a comprehensive omics analysis, with the help of transactivation assays and microscopy, we presented solid evidence on ABI5’s role in influencing ABI3’s transcriptional regulation, and how they collectively orchestrate LB dynamics during seed development and germination.

## Materials and methods

### Plant materials and growth conditions

Clustered Regularly Interspaced Short Palindromic Repeats (CRISPR)/CRISPR-associated protein 9 (Cas9) was used to generate *abi5* mutants in Arabidopsis (Wang et al., 2015). Floral dipping on *Agrobacterium tumefaciens* strain GV3101 was used to transfect ABI3-YPet-His-FLAG (hereafter ABI3YHF) transgenic lines (Clough & Bent, 1998). Selection of the transgenic lines was carried out on 1X LS medium containing 25 µg/mL Hygromycin B (GoldBio, USA, Cat # H-270-1). Columbia (Col-0) wild-type (WT), *Atabi3-6*, and ABI3YHF transgenic lines (WT and *Atabi5* background) were surface sterilized and plated on 1X Linsmaier & Skoog (LS) medium (Caisson Labs, Smithfield, UT, USA, Cat # LSP03) containing 0.7% (w/v) plant agar (PhytoTech Labs, Lenexa, KS, USA, Cat # A111). Seeds were cold stratified for 3 days at 4°C and grown on plates for 2 to 4 weeks, then transferred to soil and grown under the “long-day” condition (16 h light at 23°C / 8 h dark at 18°C).

### Plasmid construction

To produce *abi5* mutant lines, we performed CRISPR/Cas-based genome editing systems (Wang et al., 2015) .Briefly, two pairs of single guide RNAs (sgRNAs) were designed to target regions in the coding sequence of ABI5, corresponding to the N-terminal half of the protein. The targeted regions included a 379 base pair (bp) segment for the first pair and 127 bp for the second pair. The sgRNAs were amplified and subsequently cloned into pHEE401E vectors via Golden Gate assembly. ABI3YHF recombinant DNA was constructed and tagged to the C-terminal end of ABI3 using the recombineering-based gene tagging system (Alonso & Stepanova, 2015; Song, Huang, et al., 2016) and has transformed into bacmid construct. Primers used for generating CRISPR/Cas9-based *abi5* mutants are listed in Table S1.

### Germination assay

Seeds from WT and ABI3YHF transgenic lines (WT and *abi5* background) were surface sterilized and stratified at 4°C in dark for 3 days, then plated on 1X LS medium with mock, 1μM, or 3μM ABA under “long-day” condition for 10 days (Zhou et al., 2015).

### ChIP-seq library preparation

Seeds from WT, ABI3YHF/WT and ABI3YHF/*abi5* transgenic lines which weighted 0.5-1 grams were first sterilized and grown on 1X LS medium for 1 day under “long-day” condition. An additional six hours of 5μM ABA treatment was added before sample collection. Library preparation was carried out as previously described by Song et al. (Song, Koga, et al., 2016). Briefly, samples were crosslinked with 1% formaldehyde, and the nuclei were isolated. Isolated nuclei were lysed and sonicated for a total of 25 cycles of 30s ON and 90s OFF at HIGH setting in a Bioruptor Plus (Diagenode, Denville, NJ, USA). The fragmented chromatin was diluted with ChIP Dilution Buffer and were incubated overnight with FLAG antibody (1:100 v/v dilution; Sigma-Aldrich, St. Louis, MO, USA, Cat # F1804) that were bound to Dynabeads protein G (Thermo Fisher Scientific, Waltham, MA, USA, Cat # 10003D). DNA was then reverse crosslinked at 55°C overnight with an addition of 100μg/mL proteinase K before the phenol:chloroform:IAA (25:24:1 v/v, pH 8.0) extraction. Precipitated DNA was double-size selected using 0.68 vol. and 0.32 vol. SpeedBeads magnetic carboxylate (Sigma-Aldrich, St. Louis, MO, USA, Cat # GE45152105050250). Subsequently, libraries were constructed using the NEBNext® Ultra™ II DNA Library Prep Kit for Illumina® (NEB, Ipswich, MA, USA, Cat # E7645S) and sequenced on Illumina NovaSeq 6000.

### ChIP-seq analysis

Reads obtained from ChIP-Seq were analyzed with FASTQC v0.12.1 (Andrews, 2010). Low-quality reads and adaptors were trimmed by fastp v0.24.0 with default settings (Chen et al., 2018). Resulting reads were aligned to the Arabidopsis genome (TAIR10; Lamesch et al., 2012) by Bowtie 2 v2.5.4 with default settings (Langmead & Salzberg, 2012). SAM files produced were then converted to BAM files, sorted, indexed, and technical replicates were merged into one BAM file using samtools v1.9 (Li et al., 2009). Peak-calling was performed using WT control with MACS 2 v2.2.9.1 (Zhang et al., 2008). Arguments “-f BAMPE -g 120000000 -- qvalue 0.1 --call-summits” were used for specifying the input BAM files were pair-end, setting the effective genome size to 120,000,000 bp, setting the q-value threshold of 0.1 and identifying the peak summits respectively. Quality of each of the replicates were accessed through correlation and principal component analysis (PCA) plots. In short, reads were extracted from narrowPeak files and counted using RSubread v2.24.0 (Liao et al., 2013). Peaks having fewer than one total read were filtered out. The filtered counts were constructed into datasets and were normalized and variance-stabilized using median-ratio method and variance-stabilizing transformation via DESeq2 v1.50.2 (Love et al., 2014). Counts were then logarithmically transformed and used for analysis in correlation via Pearson correlation method and PCA. The correlation heatmap was drawn with pheatmap v1.0.13 (Kolde, 2025) and “RdYlBu” was selected from RColorBrewer v1.1.3 (Neuwirth, 2022). Replicates used for downstream analysis were combined with IDR v2.0.4.2 using the argument “--rank q.value --peak-merge-method min” (Li et al., 2011) and were further combined into 1 using “intersection” function from bedtools v2.31.1 with default settings (Quinlan & Hall, 2010). For quantitative analysis, csaw v1.44 was used as described by Lun and Smith (Lun & Smyth, 2016). Concisely, merged BAM files were used as input files. Reads were counted in a genome-wide sliding window with 10bp width and fragment length of 150bp. Duplicated reads and reads with a mapping quality score below 20 were excluded. Windows were retained with counts per million greater than 1 in at least 2 samples. Normalization factors were estimated from read counts in 10-kilo base pair (kb) genomic bins using the trimmed mean of M-values (TMM) method and were applied to the filtered windows. Differential binding between ABI3YHF/WT and ABI3YHF/*abi5* were accessed using edgeR v4.8.2 (Robinson et al., 2010). The filtered, TMM-normalized counts were modeled using a negative-binomial generalized linear model. Biological dispersion was estimated from the mean–dispersion relationship, and quasi-likelihood model fitting was subsequently used to account for uncertainty in dispersion estimates. Differential binding was tested using quasi-likelihood F-tests. WT control was additionally used for filtering, retaining windows that had an enrichment greater than threefold enrichment compared to WT. Adjacent windows which were within 150 bp were merged into regions with a maximum width of 500 bp.

Peaks identified from both qualitative and quantitative analysis were then annotated through ChIPseeker v1.46.1 (Yu et al., 2015). Pie charts for demonstrating ABI3 binding preference were created through obtaining annotated peak location and plotted through ggplot2 v4.0.2 (Wickham, 2010). Peaks obtained from qualitative analysis were compared against Tian et al.’s findings (Tian et al., 2020) through “intersection” function using bedtools v2.31.1 with default settings (Quinlan & Hall, 2010; Tian et al., 2020).Venn diagram was then plotted through eulerr v8.3.0 (Larsson, 2024). Histograms on the q-values from narrowPeak files and were plotted with ggplot2 v1.8.2 (Wickham, 2010). Heatmap and volcano plots were made to demonstrate annotated peaks that contained ABI3 differential binding through ComplexHeatmap v2.26.1 and ggplot2 v4.0.2 respectively (Gu et al., 2016; Wickham, 2010).

Annotated peaks were further analyzed through HOMER v4.11 for identification of binding DNA motifs (Heinz et al., 2010). DNA motifs were then clustered using motifStack v1.54 for generating clustered DNA motif trees and clouds (Ou et al., 2026). Gene ontology (GO) enrichment analysis is performed with clusterProfiler v4.18.4 with org.At.tair.db v3.22.0 as Arabidopsis genome database (Carlson, 2025; Yu et al., 2012). Transcriptomic data of *abi3-6* dry seeds were used to investigate the relationship between ABI3 binding and transcriptomic activity (Artur et al., 2026). Upset plot was plotted using ComplexHeatmap v2.26.1 (Gu et al., 2016).

Violin plot on the magnitude of changes in ABI3 binding strength and corresponding statistical analysis was performed with a one-way ANOVA test followed by Tukey’s multiple comparison in GraphPad Prism v10.3.0 (GraphPad Software, Boston, MA, USA).

Tiled data files (TDF) were generated from BAM files using igvtools v2.17.3 (Thorvaldsdottir et al., 2013). Parameters used for precomputing the maximum zoom level at 5 and setting the window size at 10 bp were “--includeDuplicates -w 10 -z 5” (Thorvaldsdottir et al., 2013). TDF files were loaded to Integrative Genomics Viewer v2.19.2 with Araport11 annotations were used to visualize read distribution in the genome (Cheng et al., 2017).

### RNA-extraction and qPCR

Seeds from Col-0 WT, ABI3YHF/WT and ABI3YHF/*abi5* transgenic lines which weighed 0.5-1 grams were first sterilized and grown on 1X LS medium for 1 day under “long-day” condition. 5μM of ABA was added for an additional 6 hours prior to sample collection. Total RNA is extracted using Plant/Fungi Total RNA Purification Kit (Norgen Biotek, Thorold, ON, Canada, Cat # 31350). Extracted RNA were treated with DNase using Invitrogen^TM^ TURBO DNA-free^TM^ Kit (Invitrogen, Carlsbad, CA, USA, CAT # AM1907) and were reverse transcribed to cDNA using Thermo Scientific^TM^ Maxima H Minus Reverse Transcriptase (Thermo Fisher Scientific, Waltham, MA, USA, Cat # EP0752). Quantitative PCR (qPCR) was conducted using PowerUp^TM^ SYBR^TM^ Green Master Mix (Thermo Fisher Scientific, Waltham, MA, USA, Cat # A25742). qPCR primers are listed in Table S2. qPCR primers, which were previously published, for SEIPIN1 (Grace et al., 2023), OLEO1 (Traver & Bartel, 2023), OLEO2 (Traver & Bartel, 2023), and OLEO5 (Zhao et al., 2021) were used.

## Results

### abi5 mutants in ABI3YHF transgenic lines exhibited ABA insensitivity during germination

ABI3YHF transgenic lines were created through transformation-competent bacterial artificial chromosome to preserve the regulatory context of ABI3 in *Arabidopsis thaliana* (*A. thaliana*) (Song, Huang, et al., 2016). To generate *abi5* mutants in ABI3YHF transgenic lines, we used CRISPR-Cas9 systems to mutate ABI5 coding sequences. Three independent mutant lines, namely ABI3YHF-1/*abi5-11*, ABI3YHF-1/*abi5-12* and ABI3YHF-2/*abi5-13* was created. Sequencing results revealed indel mutations leading to premature stop codons upstream of bZIP domain (Fig. 1A). *ABI3* expression in *abi3-6* mutants was completely abolished, whereas ABI3YHF transgenic lines (WT and *abi5* mutant backgrounds) contained nearly 2.5-fold higher than WT (Fig. 1B). Meanwhile, *ABI5* expression was abolished in *abi3* and ABI3YHF/*abi5* mutants while ABI3YHF/WT contained similar expression level as WT (Fig. 1B).

**Figure 1.**
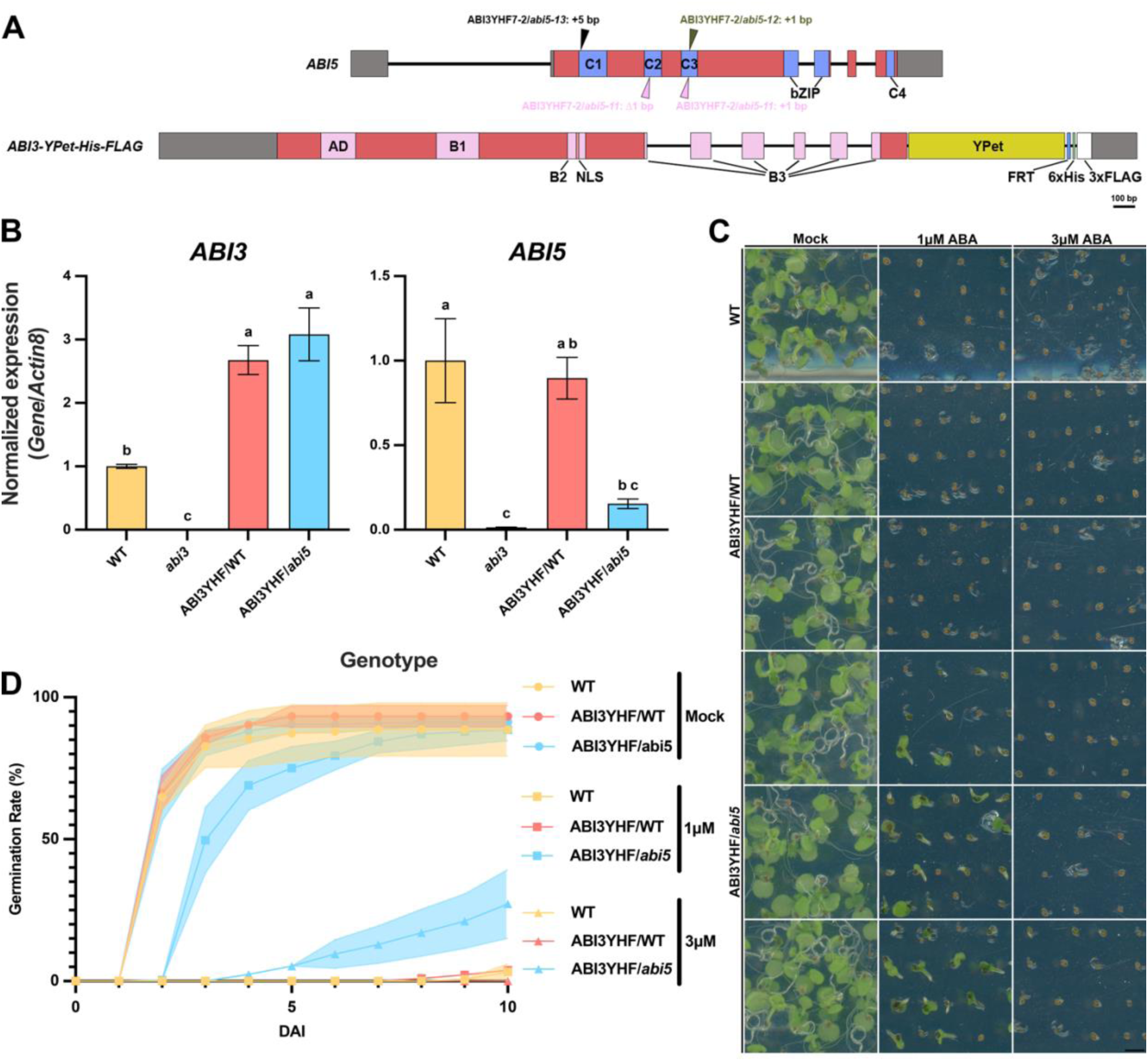
*abi5* mutants in ABI3YHF transgenic lines exhibit reduced sensitivity to ABA (A) Genomic ABI5 and ABI3YHF sequence is presented, with grey areas representing untranslated regions, red areas representing exons, and black lines representing introns. Blue boxes represent domains present in ABI5 protein. Pink boxes represent domains present in ABI3 protein. Salmon box represents NLS. Yellow, sky-blue, green, and white box corresponds to YPet, flippase recognition target (FRT), histidine (His), and FLAG DNA sequences. Arrows represent location of mutation, with black, pink, and dark yellow representing independent lines *abi5-11*, *abi5-12*, and *abi5-13*. Scale bar = 100bp. (B) Expression level of *ABI3* and *ABI5* in WT (Gold), *abi3* (white), ABI3YHF/WT (salmon) and ABI3YHF/*abi5* (sky-blue). (C, D) Germination rate of 10 DAI ABI3YHF transgenic lines of ABI3YHF/WT and ABI3YHF/*abi5* mutants with WT as control on plates treated with none, 1μM, or 3μM ABA. Yellow, salmon, and sky-blue color represents WT, ABI3YHF/WT, and ABI3YHF/*abi5* germination rate respectively. Circle labeled lines refers to germination rate under mock treatment; rectangle labeled lines refers to germination rate under 1μM treatment; triangle labeled lines refers to germination rate under 3μM treatment. Scale bar = 5mm.

To examine if *abi5* mutants are ABA insensitive, we tested the germination rate of WT and ABI3YHF transgenic lines (WT and *abi5* mutant backgrounds) with or without 1μM or 3μM ABA. While all lines exhibited similar germination rate in mock treatment, germination rate of WT and ABI3YHF/WT transgenic lines was inhibited with 1μM or 3μM ABA treatments (Fig. 1C, 1D). Meanwhile, ABI3YHF/*abi5* transgenic lines had delayed germination under 1μM or 3μM ABA treatments (Fig. 1C, 1D). In 1μM ABA treatment, ABI3YHF/*abi5* had median germination time (T50) of 2DAI while germination rate if ABI3YHF/*abi5* were around 25% at 10DAI in 3μM ABA treatment, unlike mock treatment when T50 was 1DAI (Fig. 1D). These results confirmed the CRISPR *abi5* lines reduced the sensitivity to ABA. WT and ABI3YHF/WT seeds failed to germinate under 1μM and 3μM ABA treatment. Interestingly, *abi5-8* has a rate of cotyledon greening around 70% (Zhou et al., 2015). The discrepancy between ABI3YHF/*abi5* mutants and *abi5-8* might be the overexpression of ABI3 proteins, as ABI3YHF reporter line was introduced in WT background instead of *abi3* mutant background (Fig. 1B).

### ABI3 ChIP revealed reduced binding strength in ABI3-induced genes in abi5 mutants

ABI5 has been known to be an interactor of ABI3 (Nakamura et al., 2001). Yet, exact evidence on how ABI3’s binding preference is influenced by ABI5 is not known. To examine the binding patterns of ABI3, we performed ChIP-seq on ABI3YHF/WT and ABI3YHF/*abi5*. Using WT as control, ABI3/WT and ABI3/*abi5* transgenic lines formed two distinct clusters, with one of the ABI3/WT samples being the outlier (Fig. S2A, 2B). Annotation of bound peaks revealed that the binding location of ABI3 did not change. where there were around 75% of peaks located at the proximal promoter (within 1 kb from transcription start site (TSS)) (Fig. 2A) (Table S3) (Morales-Cruz et al., 2026). There were 5420 and 7388 peaks discovered in WT background and *abi5* mutant background respectively, with 4820 peaks that were shared between both backgrounds (Fig. 2B). Intriguingly, ABI3YHF/*abi5* mutants revealed higher number of peaks compared to ABI3YHF/WT, yet the q-values of ABI3YHF/*abi5* mutants was smaller than that of ABI3YHF/WT (Fig. S2). Comparing with the latest ChIP experiment on ABI3 TF, our results discovered approximately 83% of peaks that were also annotated in literature, along with many new binding sites (Fig. 2B).

**Figure 2.**
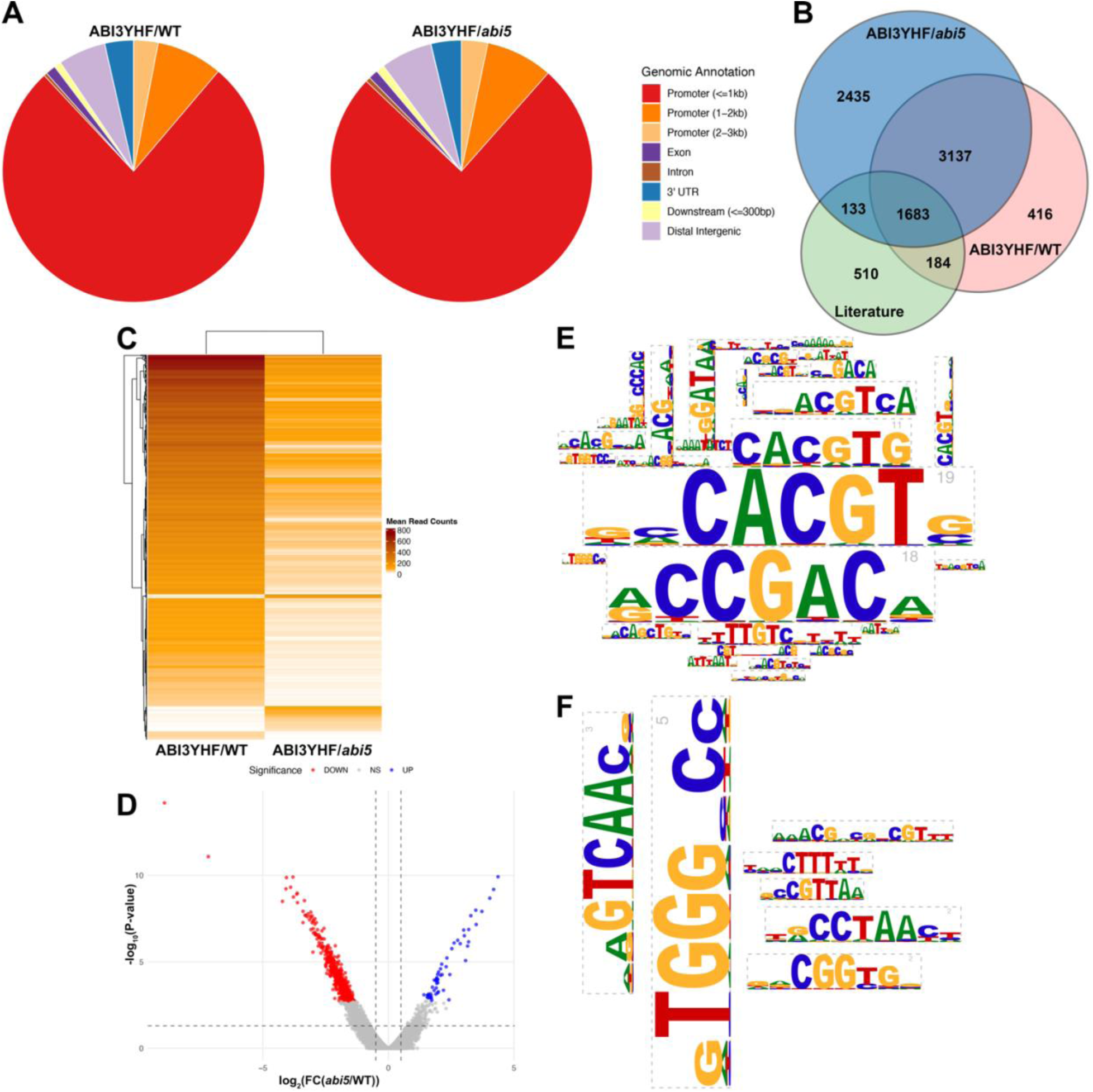
ABI3 binding strength and preferences are altered in *abi5* mutant (A) Distribution on binding preferences of ABI3YHF/WT and ABI3YHF/*abi5*. (B) Venn diagram on peaks found from ChIP-seq of ABI3YHF/WT and ABI3YHF/*abi5* mutants, and ChIP-ChIP results of ABI3 from Tian et al.’s results () (C) Heatmap showing the read counts of the peaks identified by ABI3YHF/WT and ABI3YHF/*abi5* mutants. (D) Volcano plot shows the number of peaks having increased (blue) or decreased (red) ABI3 binding in *abi5* mutant background compared to WT background. (E, F) Clustered enriched DNA motifs bound by (E) decreased and (F) increased binding of ABI3YHF/*abi5* mutants compared to ABI3YHF/WT

We further examined the DNA binding patterns of ABI3YHF/WT and ABI3YHF/*abi5* mutants using HOMER (Heinz et al., 2010) for DNA motif identification and motifStack for motif clustering (Ou et al., 2026). In both ABI3YHF/WT and ABI3YHF/*abi5* mutants, the G-Box motif and its core sequence “ACGT” were found, showing that despite the presence of ABI5 TF, ABI3 can still bind to G-Box motifs (Fig. S4 - S6). Notably, the clustering of G-Box related motifs was less enriched in ABI3YHF/*abi5* mutants compared to ABI3YHF/WT, hinting a reduced binding towards G-Box motifs (Fig. S4 - S6). Interestingly, in both ABI3YHF/WT and ABI3YHF/*abi5* mutants, G-Box motifs were not the most enriched motif being clustered (Fig. S4 - S6). Motifs of “TTGAC[C/T]” were highly enriched (Fig. S4 - S6). This motif corresponds to the WRKY TF family (Ciolkowski et al., 2008). This indicates that the binding ability of ABI3 towards the WRKY motif is hindered by ABI5. Other than that, the binding motif GC-rich “GTGCCG” and “[T/A]AAAG” were highly enriched in both ABI3YHF/WT and ABI3YHF/*abi5* mutants (Fig. S5, S6B). These motifs correspond to the APETALA2/ETHYLENE-RESPONSIVE ELEMENT BINDING PROTEIN (AP2/EREBP) and Cys2/Cys2 DNA-binding with one finger (C2C2Dof) zinc finger TF family respectively (Hao et al., 1998; Vicente-Carbajosa & Carbonero, 2005). However, the enrichment was more significant in *abi5* mutant backgrounds compared to WT backgrounds (Fig. S4 - S6). This may imply that the absence of ABI5 TF opens up ABI3’s binding capacity towards these motifs.

To examine if the binding strength of ABI3 has been shifted, we performed quantitative analysis using csaw (Lun & Smyth, 2016). Comparing with the qualitative analytic pipeline, quantitative analysis revealed approximately 70% peaks that were also found in the qualitative analytic pipeline (Fig. S7A). Nonetheless, the peak width was significantly smaller than the qualitative analytic pipeline, with an average of around 1kb in the qualitative analytic pipeline and below 500bp in the csaw quantitative analytic pipeline (Fig. S7B, S7C). This provides higher precision of the location of peak. Overall speaking, ABI3YHF/WT accounted for a higher number of read counts compared to ABI3YHF/*abi5* mutants, with two groups having higher number of read counts in ABI3YHF/*abi5* (Fig. 2C). There were 777 decreased ABI3 binding peaks and 79 increased ABI3 binding peaks in *abi5* mutant background (Fig. 2D). DNA motif enrichment analysis revealed that peaks with decreased ABI3 binding contains motifs that were highly related to G-Box whereas peaks with increased ABi3 binding contains motifs related to TEOSINTE BRANCHED 1, CYCLOIDEA, and PROLIFERATING CELL FACTOR (TCP), WRKY, and AP2/EREBP TF families (Fig. 2E, 2F, S8).

### ABI3 regulates lipid body related genes, and ABI5 fine tunes ABI3 binding strength

We further compiled ABI3 binding information with publicly available transcriptomic dataset of *abi3-6* dry seeds to investigate the transcriptomic significance of changes in ABI3 binding’s binding strength due to ABI5 interaction (Artur et al., 2026). The group with most genes (332 genes) were genes that were induced by ABI3 and had decreased ABI3 binding in *abi5* mutant (Fig. 3A). The group with second most genes (195 genes) were genes that were repressed by ABI3 and had decreased ABI3 binding in *abi5* mutant (Fig. 3A). The remaining two groups were related to increased ABI3 binding in *abi5* mutant, which only accounted for a total of 40 genes (Fig. 3A) (Table S4). When further quantifying the magnitude of ABI3 binding in all 4 groups, ABI3 induced genes revealed the greatest change in ABI3 binding in *abi5* mutant background, showing ABI5 exerts larger influence on ABI3’s binding in inducing gene expressions (Fig. 3B).

**Figure 3.**
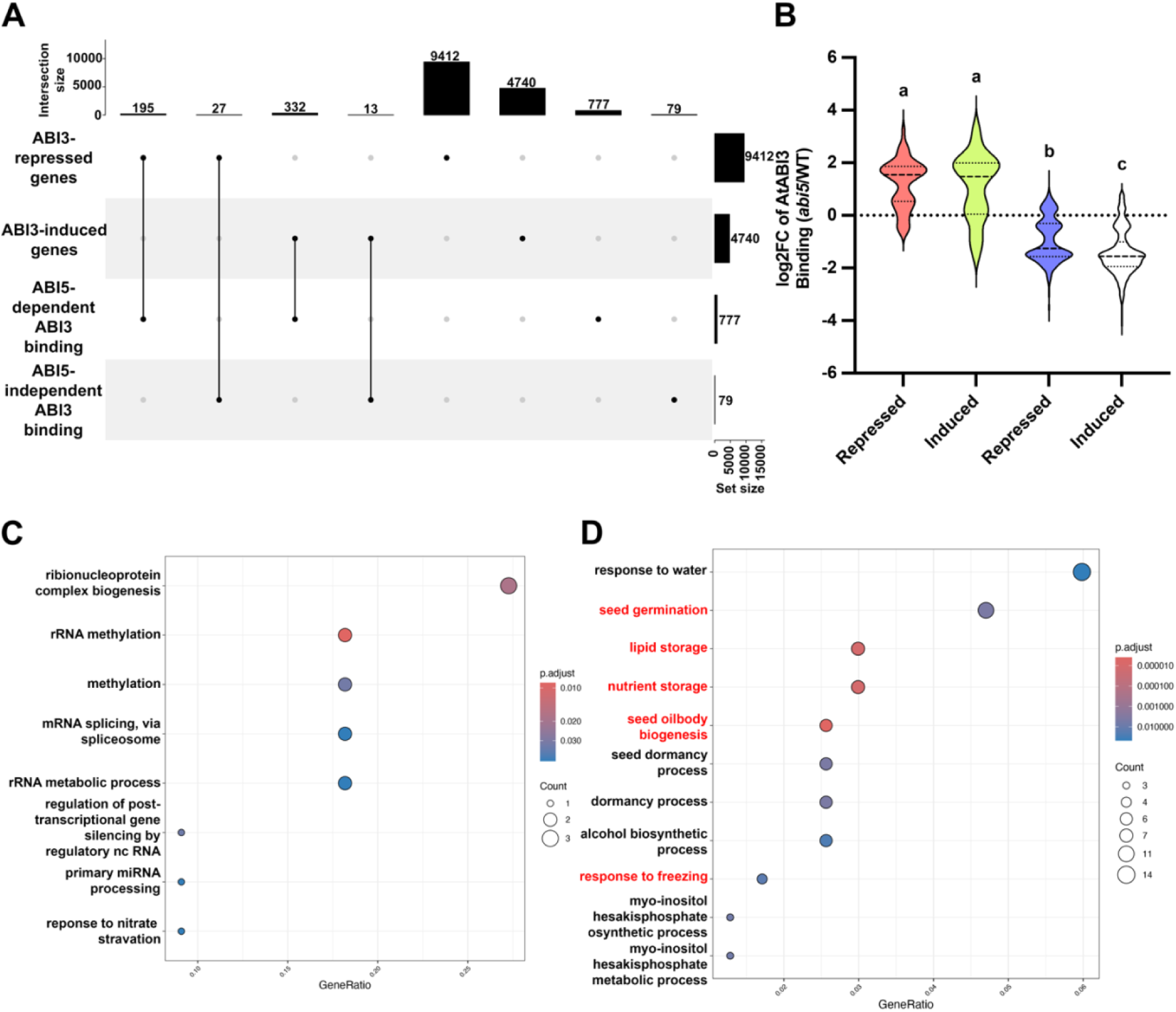
ABI5 has higher influence on ABI3’s binding towards ABI3 induced genes (A) Upset plot showing genes that are shared between increased or decreased binding of ABI3 in *abi5* mutant background and increased or decreased expression in *abi3* mutant. (B) Violin plot of magnitude in changes of ABI3 binding in *abi5* mutant in ABI3-induced or -reduced genes. (C, D) GO enrichment analysis in biological processes in (C) ABI5-dependent ABI3 repressed genes and (D) ABI5-dependent ABI3 induced genes. Processes related to LB dynamics were highlighted in red.

To investigate what biological processes is governed by both ABI3 and ABI5, we performed GO enrichment analysis on genes that were downregulated in *abi3* mutant and had reduced ABI3 binding in *abi5* mutants and genes that were upregulated in *abi3* mutants and had reduced ABI3 binding in *abi5* mutants. Enrichment in molecular functions in genes that were upregulated in *abi3* mutants and had reduced binding ABI3 binding in *abi5* mutants showed processes related to transcriptional and translational regulation (Fig. 3C). On the other hand, enrichment analysis on genes that were downregulated in *abi3* mutants and had reduced ABI3 binding in *abi5* mutants and genes that were upregulated in *abi3* mutants revealed process related to seed development and germination, for instance, seed germination, lipid storage, nutrient storage, and seed oil body biogenesis (Fig. 3D).

We focused on processes related to lipid storage and further inspected genes related to such processes (Fig. 4A). Genes related to lipid body biogenesis, including oleosins and SEIPIN1, lipid body associated proteins, such as OBAP1A and OBAP2A, and lipid body fusion protein LDPS were found (Fig. 4B) (Table S5) (Guzha et al., 2023). We further examined transcript levels of these genes to validate if they are collectively regulated by ABI3 and ABI5. *abi3* mutants showed absence of these genes, showing that ABI3 is necessary for the presence of these proteins (Fig. 4C). ABI3YHF/WT transgenic lines contained expression levels 1.5 to 2.5 times higher than WT, with the exception of *LDPS* (Fig. 4C). For ABI3YHF/*abi5*, *LDPS*, *OBAP2A*, and *OLE5* had expression levels halved compared to WT (Fig. 4C). Although *OBAP1A*, *OBAP2C*, *SEIPIN1*, *OLE1*, and *OLE2* contain comparable expression levels with WT, its expression level was at least halved compared to ABI3YHF/WT (Fig. 4C). This further supports that ABI5 promotes binding of ABI3 for transcriptional activation of these genes (Fig. 4C).

**Figure 4.**
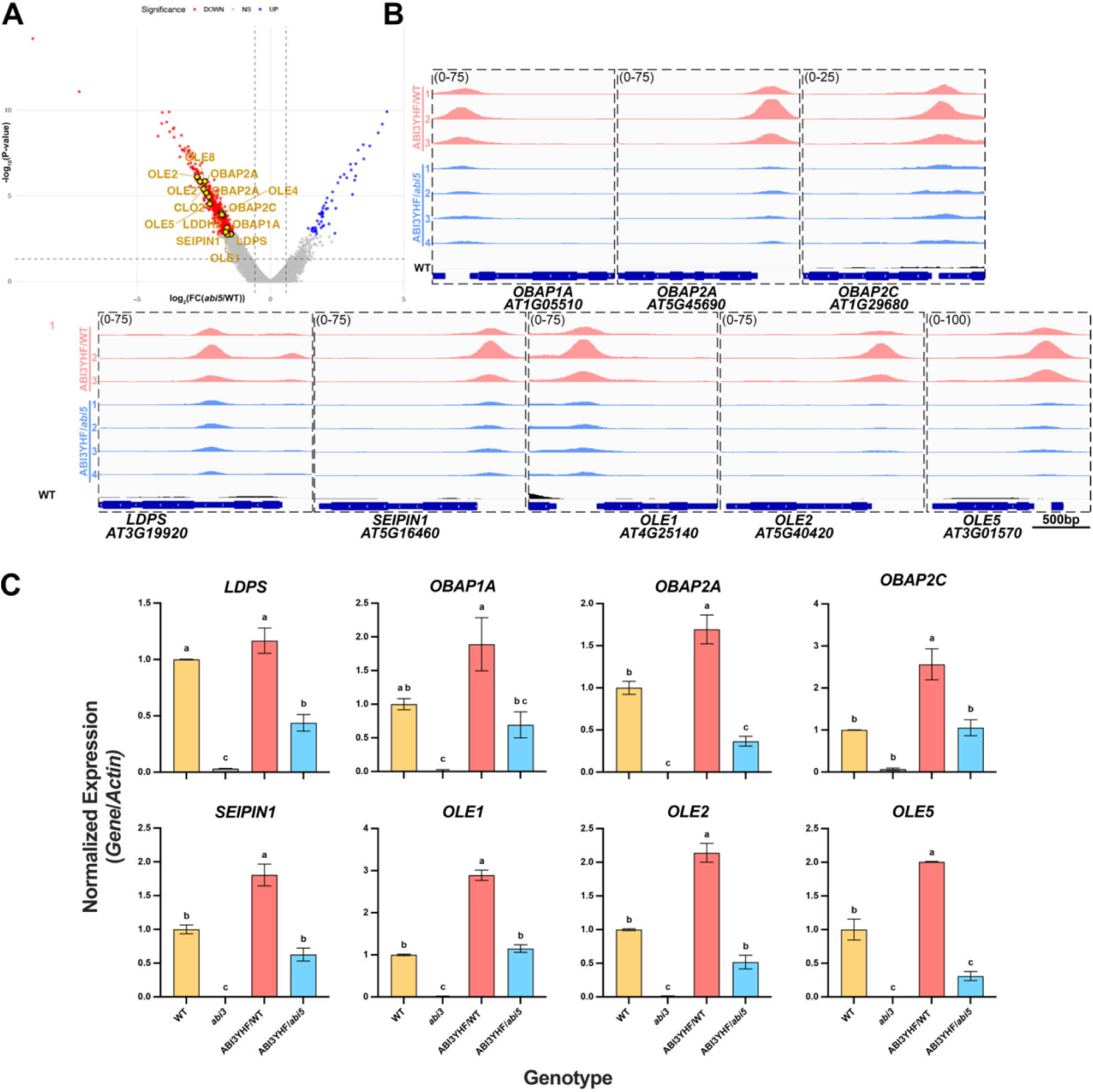
Multiple lipid body genes are coordinately regulated by ABI3 and ABI5. (A) Volcano plot shows the number of peaks having increased (blue) or decreased (red) ABI3 binding in *abi5* mutant background compared to WT background with gold dots highlighting genes related to LB dynamics. (B) Integrative Genomics Viewer (IGV) showing peaks of ABI3YHF/WT (salmon) and ABI3YHF/*abi5* mutants (sky-blue) in lipid body related proteins. Scale bar = 500bp. (C) Expression level of LB related genes in WT (Gold), *abi3* (white), ABI3YHF/WT (salmon) and ABI3YHF/*abi5* (sky-blue).

**Figure 5.**
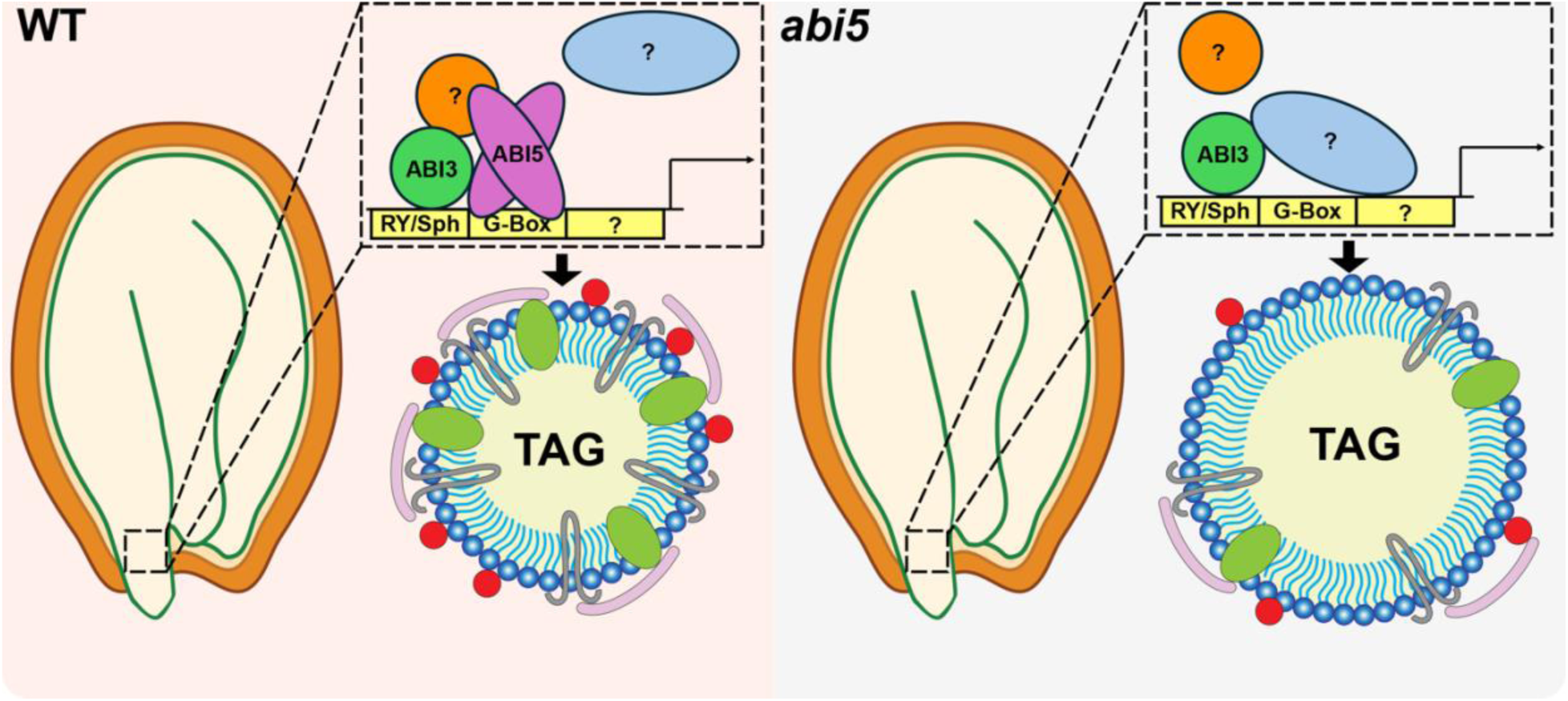
Hypothesized model of ABI5-dependent ABI3 binding in regulating lipid body dynamics In WT germinating seeds, ABI5 binds to ABI3 to enhance the binding towards G-Box elements. This leads to the upregulation of LB related genes and respective proteins depositing on to LB. As proteins such as oleosins are expressed, it allows the LB to maintain small volumes by preventing LB fusion. On the other hand, in *abi5* mutants, the absence of ABI5 does not abolish ABI3’s binding towards G-Box motifs, as other TFs such as members from C2C2dof Zinc finger family binds to CREs near G-Box motifs. Yet, ABI3’s binding has weakened. This leads to lower expression of LB related genes and respective proteins. Thus, with lower proteins like oleosins deposited in LB, LB dynamics is disrupted, leading to enlarged LB.

## Discussion

Transcriptomic regulations during phase transition are extremely complicated, as numerous gene expressions change to support diverse changes in both cellular, tissue, and physiological levels. For example, during seed development, LEC1, one of the four master regulators and the earliest pioneer factor dictating transcriptional regulation during seed development, is known to act in combinatorial interactive manner with different TFs such as ABI3, and bZIP67 and ABSCISIC ACID-RESPONSIVE ELEMENT BINDING PROTEIN 3 (AREB3) (Jo et al., 2020). The different combinations not only operate in a differential temporal manner but also bind to different cis-regulatory element (CRE) regions, governing different biological processes at different time points during seed germination (Jo et al., 2020). ABI3 is well known to be one of the master regulators during seed development in which it controls multiple biological processes such as nutrient storage. Interestingly, although it is well documented that it regulates diverse gene expressions, the details of fine tuning or mechanisms of transcriptional regulatory mechanisms from ABI3 is not well known. Here, we revealed one of the fine-tuning mechanisms on ABI3’s transcriptional regulation on lipid bodies with the assistance of ABI5.

### ABI3 has a high bias towards G-Box motif regardless of ABI5 compared to RY/Sph motif

ABI3 has been well known to bind to both RY/Sph and G-Box motifs (Ezcurra et al., 2000; Tian et al., 2020). The RY/Sph element is bound by the B3 domain whereas the G-Box motif is bound by the B2 domain, where the B2 domain also possesses nuclear localization signal (NLS) (Ezcurra et al., 2000). Intriguingly, the binding ability towards G-Box motifs are lineage specific (Ezcurra et al., 2000; Sakata et al., 2010; Tian et al., 2020). Members of bZIP TFs interacting with ABI3 are also known to bind to G-Box motifs through the bZIP domain (Izawa et al., 1993). We hypothesize that the interaction between ABI3 and bZIP TFs further promote the binding towards G-Box motifs. Indeed, even in the absence of ABI5, ABI3 still significantly bound towards G-Box motifs (Fig. S4 - S6). Interestingly, ABI3YHF/WT and ABI3YHF/*abi5* mutants revealed minimal binding towards the RY/Sph motif (Fig. S4 - S6). This may be due to the weak binding ability of ABI3 towards the RY/Sph motif compared to other members in the same family, FUS3 and LEC2 (Jia et al., 2021). It has been demonstrated that the conserved B3 domain has slight divergence between the B3 domain, where ABI3 lacks the crucial K64, K66 and S69, leading to weakened B3 binding ability (Jia et al., 2021). These would explain the exceptionally low binding events on RY/Sph elements.

The binding ability of ABI3 towards G-box motifs are divergent throughout land plants. While bZIP TFs further promote the binding preference towards G-Box motifs, ABI3 in higher plants already possesses the ability to bind to G-Box. This is reflected in our results, where the absence of ABI5 also possessed a relatively high enrichment in binding towards G-Box motifs (Fig. S5, S6B), although the possibility of ABI3 interacting with or in close proximity with other bZIP or basic helix-loop-helix (bHLH) TFs must not be ignored, for instance, interactions with PHYTOCHROME INTERACTING FACTOR 1 (PIF1 or otherwise known as PIL5) has been previously reported (Park et al., 2011). ABI3 and PIF1 collectively regulate *SOMNUS* (*SOM*) during imbibition, where ABI3 and PIF1 bind to CRE containing proximal RY/Sph elements and E-box motifs, while solely ABI3 is sufficient to promote *SOM* expression during seed maturation (Park et al., 2011). SOM in turn promotes ABA biosynthesis and suppresses gibberellin (GA) biosynthesis (Park et al., 2011). While direct evidence of interaction between ABI3 and other bHLH and bZIP TFs is lacking, bHLH and bZIP TFs have high tendency in binding G-box motifs clustered with RY/Sph elements (termed as RY/G clusters) in *Brassicae napus* (Martin et al., 2008; Verdier & Thompson, 2008). Combining our results, we propose that ABI3’s binding towards G-Box gains higher impact in transcriptional activity. This may imply that bHLH and bZIP TFs localized near RY/G clusters are important for ABI3-mediated transcriptional regulations during seed development. More on ABI3’s binding property can be explored through comparing ABI3 orthologs and their respective DNA binding preferences within land plants to grasp a deeper understanding on how ABI3 binds to and regulates gene expressions.

### ABI3’s interacting partner competes with ABI5 for DNA binding

While ABI3 can still bind to other motifs besides the G-Box motif, the binding ability is enhanced under the absence of ABI5. DNA motifs which were recognized by AP2/EREBP and C2C2dof Zinc Finger TFs were bound and enhanced in *abi5* backgrounds. ETHYLENE RESONSIVE FACTOR 1 (ERF1) is one of the AP2/EREBP TFs that interact with ABI3. The ABI3-ERF1 module is believed to act as a molecular rheostat to avoid over stimulation from both ABA and auxin signalling through inhibiting ABI3 and ERF1 from binding DNA, thus inhibiting lateral root formation (Zhang et al., 2023). ABI4 is another AP2/EREBP TF that is tightly correlated to seed developmental processes and ABA signalling. While ABI3 and ABI4 do not have interaction in yeast systems (Nakamura et al., 2001), they perform synergistic regulations of genes promoting seed developmental programs. For instance, the absence of ABI4 would attenuate expressions of storage protein, *RESPONSIVE TO ABA* 18 (*RAB18*), and *ABI3* (Soderman et al., 2000). This suggests that ABI4 partially participates in the regulatory network of ABI on seed developmental genes. Regarding the binding motif of C2C2dof Zinc Finger, its binding motif position is slightly further from neighbouring RY/Sph and G-Box motif in promoters of seed storage protein genes, closer towards the TSS (Morales-Cruz et al., 2026; Vicente-Carbajosa & Carbonero, 2005). The increased enrichment of AP2/EREBP and C2C2dof Zinc Finger motifs in *abi5* mutants indicates that ABI5 has a higher priority in interacting with ABI3 and subsequent binding towards CRE regions. When ABI5 is absent, AP2/EREBP and C2C2dof Zinc Finger TFs can then bind to their respective motifs proximal to the RY/G cluster motifs. This explains even under the loss of ABI5, the seed maturation program still persists as these alternative binding sites serve as an alternative or “backup” plan to regulate seed developmental genes. Yet, the binding strength of ABI3 is not as strong as with the assistance from ABI5. This may also explain the lowered expression level of LB related genes in *abi5* mutants (Fig. 4C).

### ABI3 and ABI5 collectively regulate lipid body dynamics during seed development and germination

Lipid bodies are organelles that serve multiple functions like energy reservoir, membrane source, and stress responses in organisms (Lundquist et al., 2020). In plants, LBs are in large abundance in seeds, pollen, and fruits, mainly serving as energy reservoirs and membrane sources (Bouchnak et al., 2023). In other plant tissues, LBs’ main function lies during stress response, where the accumulation of LBs provides resources for sequestering excess lipids and membrane remodelling (Bouchnak et al., 2023).

Like oleosins, OBAPs are evolutionary conserved proteins that are present across different kingdoms (Lopez-Ribera et al., 2014). In seed plants, OBAPs are exclusively found in seeds (Di et al., 2025; Lopez-Ribera et al., 2014; Lujan et al., 2025). While OBAPs does not contain signal peptides nor long hydrophobic regions to be retained in membranes, it is localized to the surface of LBs (Lopez-Ribera et al., 2014). Mutation of OBAP1A proteins leads to lowered germination rate in *A. thaliana*, with enlarged and irregular shaped LBs and altered lipid content (Lopez-Ribera et al., 2014). On the other hand, ectopic expression of *Leymus chinensis OBAP2B* in Arabidopsis increased its salt tolerance as the germination rate of Arabidopsis increased under salt treatment (Di et al., 2025). These indicated that OBAPs function in regulating lipid body during seed development and stress responses are mostly likely conserved in seed plants. Our finding on OBAP families co-regulated by ABI3 and ABI5 further supports the idea. This further indicates ABI3 and ABI5’s involvement in regulating LB in seeds through maintaining LB structural integrity on top of oleosins.

Recent studies have analysed LDPS’s role during seed development and germination. LDPS is involved in LB fusion during seed germination, which it is speculated to either reduce packing deficiency of LBs as oleosins degrade or to slow down TAG metabolism for matching the rate of β-oxidation (Doner et al., 2025). Given the results of ABI3 and ABI5 co-regulating the transcription of LDPS, it is intriguing to evaluate the temporal expression level of LDPS protein. ABI3 expression level degrades as seed enters dormancy and germination, while ABI5 plays a role in early post-germinative stages. Such results may indicate that LDPS proteins are synthesized early during seed germination stage, yet, due to “crowding effect”, it out-competes with other structural proteins such as oleosins and OBAPs, therefore, unable to be anchored directly on the LB membrane and promote LB fusion. During germination, as oleosins and OBAPs started to degrade first, the presence of LDPS promotes LB fusion, executing its significance on governing LB dynamics.

While most of the genes displayed in this study are related to LB dynamics, it is also worth noting that there is drastic lipid metabolic activity during seed development and germination for synthesizing and metabolizing lipid nutrients respectively (Bouchnak et al., 2023). WRINKLED1 (WRI1) is one of the major TFs regulating lipid metabolism during seed germination (Kong et al., 2026). For example, WRI1 regulates FATTY ACID DESATURASE 2 (FAD2) for synthesizing linolenic acid, one of the main TAGs stored in seed (Kong et al., 2026). While ABI3’s ability to upregulate WRI1 is well known, ABI5’s ability to regulate WRI1 was found solely in oil palm (Kong et al., 2026). In our ChIP-seq results, we did not see a high binding towards WRI1, potentially due to the stage of the samples as we collected ABA-treated germinating seeds instead of developing seeds. On the other hand, DGATs are also well known to be regulated by both ABI3 and ABI5 (Kong et al., 2013; McGuire et al., 2025). While we found binding of ABI3 towards *DGAT*s’ promoters, absence of ABI5 did not significantly alter ABI3’s binding strength. Perhaps the regulation of DGAT from ABI3 and ABI5 are independent from each other, while ABI5’s regulation is wired with ABI4 (Kong et al., 2013). This further provides evidence on emphasis of ABI3 and ABI5’s role towards regulating LB dynamics instead of lipid metabolism. Nevertheless, these revealed that the regulation of LB by ABI3 and ABI5 are not solely on the synthesis of LBs during seed development, but a more complicated mechanism and physiology in governing LB dynamics.

Taken together, we have demonstrated how ABI5 promotes ABI3’s transcriptional regulation on genes related to seed development and germination through ChIP-seq analysis combining with existing RNA-seq data available and used LB as an example to demonstrate the combined transcriptional regulatory mechanism of ABI3 and ABI5. We propose that normally ABI5 interacts with ABI3. This enhances ABI3’s binding towards G-Box motifs, leading to upregulation of LB related genes and consequently deposition of LB related proteins onto LB. Expression of LB related proteins, including oleosins and OBAPs, prevent LB fusion, thus, having smaller LB. Whereas in *abi5* mutants, absence of ABI5 attenuates ABI3’s binding towards G-Box motifs. ABI3 can still bind towards G-Box motifs with potential protein interactors such as members from C2C2dof Zinc finger family. The weakened binding leads to lower expression level of LB related genes, therefore, proteomics in LB is disrupted, leading to unusual LB dynamics such as enlarged LB (Fig. 6). These findings further complemented missing information on not only how ABI3 binds to certain subsets of genes but also the delicate collective regulation from ABI3 and ABI5 on LBs. It will be interesting to see if the relationship between ABI3 and ABI5 are conserved throughout land plants since ABI3 and ABI5 are ancient proteins that exist in early diverging land plants.

## Acknowledgement

The authors thank Dr. Nambara (University of Toronto) for providing *abi3-6* mutant. The authors also thank funding support from the University of British Columbia Four Year Doctoral Fellowships (UBC 4YF) to C. C. H. H and M. A, and NSERC RGPIN-2019-05039 and CFI JELF/BCKDF 38187 to L.S.

