## Supplementary figures 1-7 and supplementary tables 1-5 for "ABSCISIC ACID INSENSITIVE 5 fine-tunes expression of lipid body genes through modulating ABSCISIC ACID INSENSITIVE 3 binding strength"

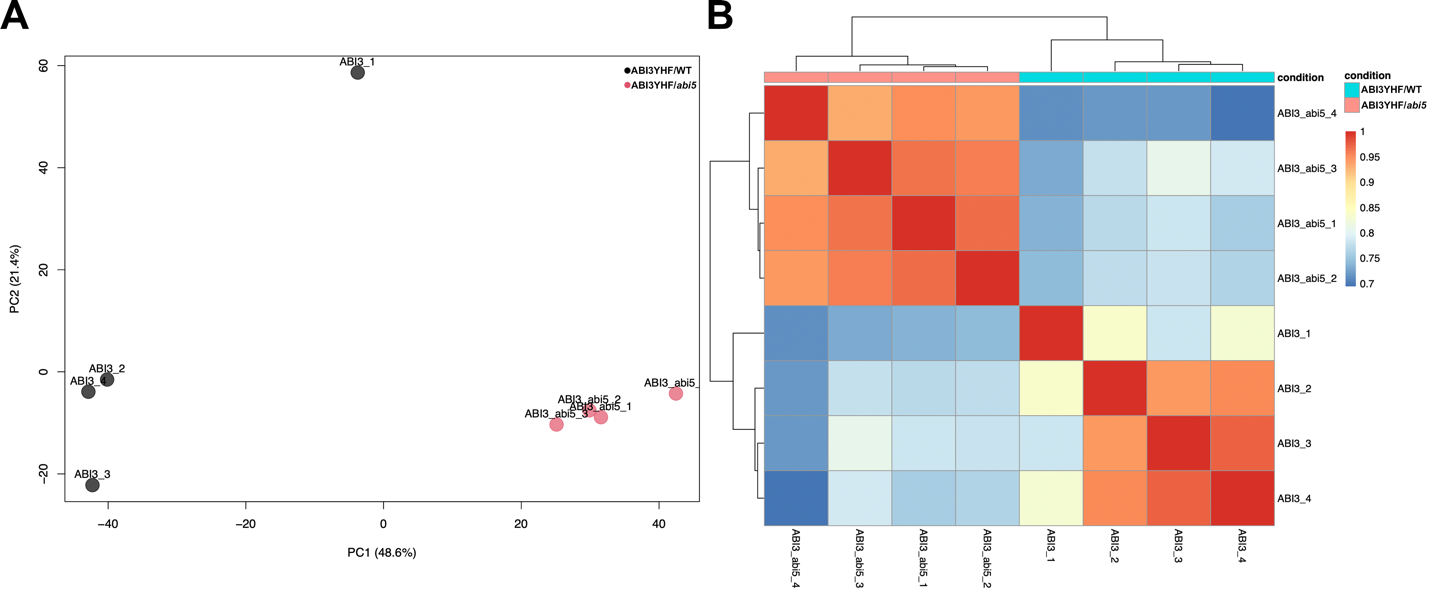

Figure S1. Quality check of ABI3 ChIP-Seq results

(A) PCA plot showing variance between each sample from ABI3 ChIP-seq results. (B) Pearson correlation heatmap on each sample from ABI3 ChIP-seq results.

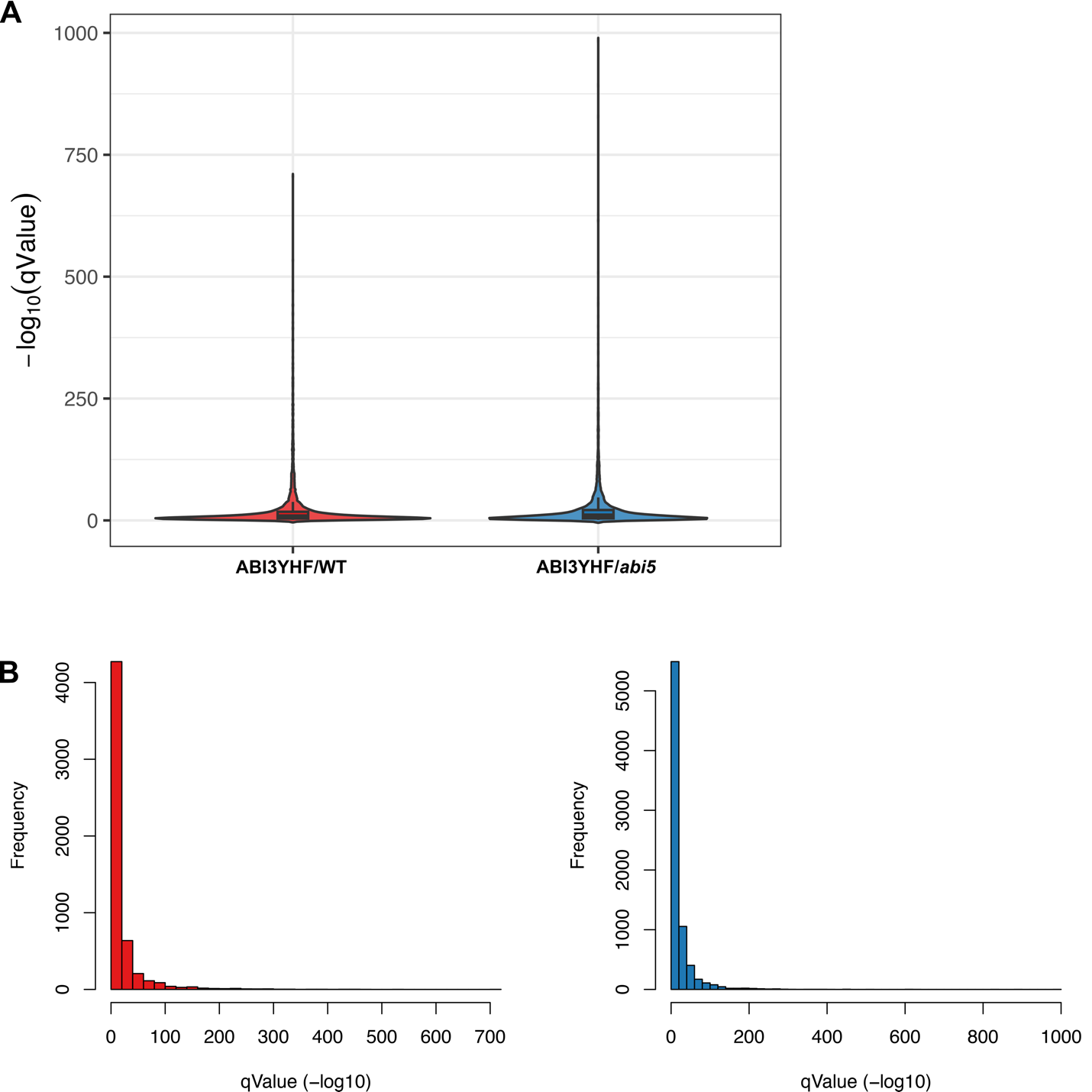

Figure S2. Number of peaks and peak quality in WT and *abi5* mutant backgrounds

(A, B) Q-values of peaks from ChIP-seq of ABI3 from WT and *abi5* mutants showing in the form of (A) violin plot and (B) histogram.

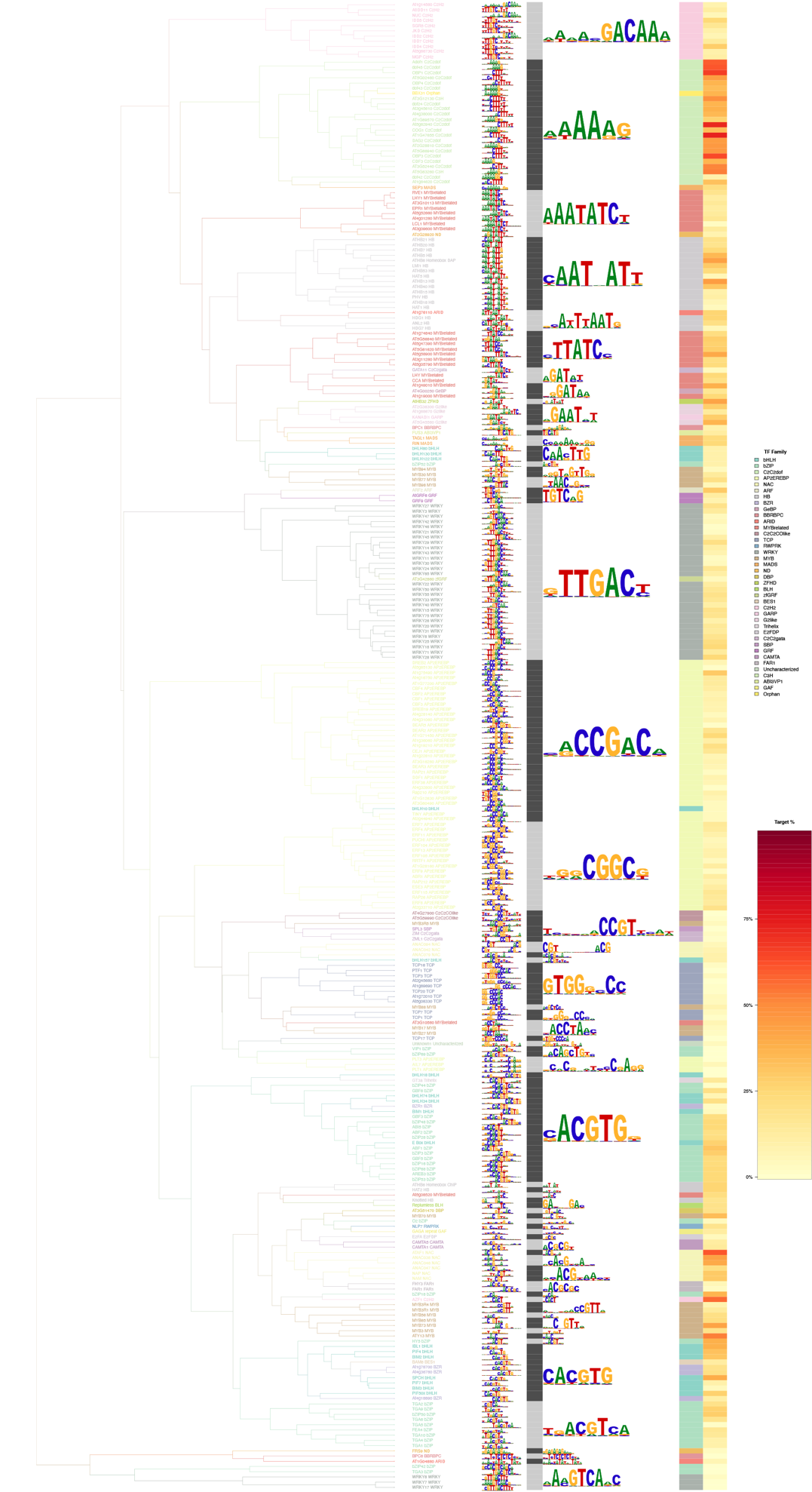

Figure S3. Motif tree on enriched motifs bound by ABI3 in WT background

Enriched motif tree found in DNA motifs from HOMER analysis in ChIP-seq ofABI3YHF/WT.

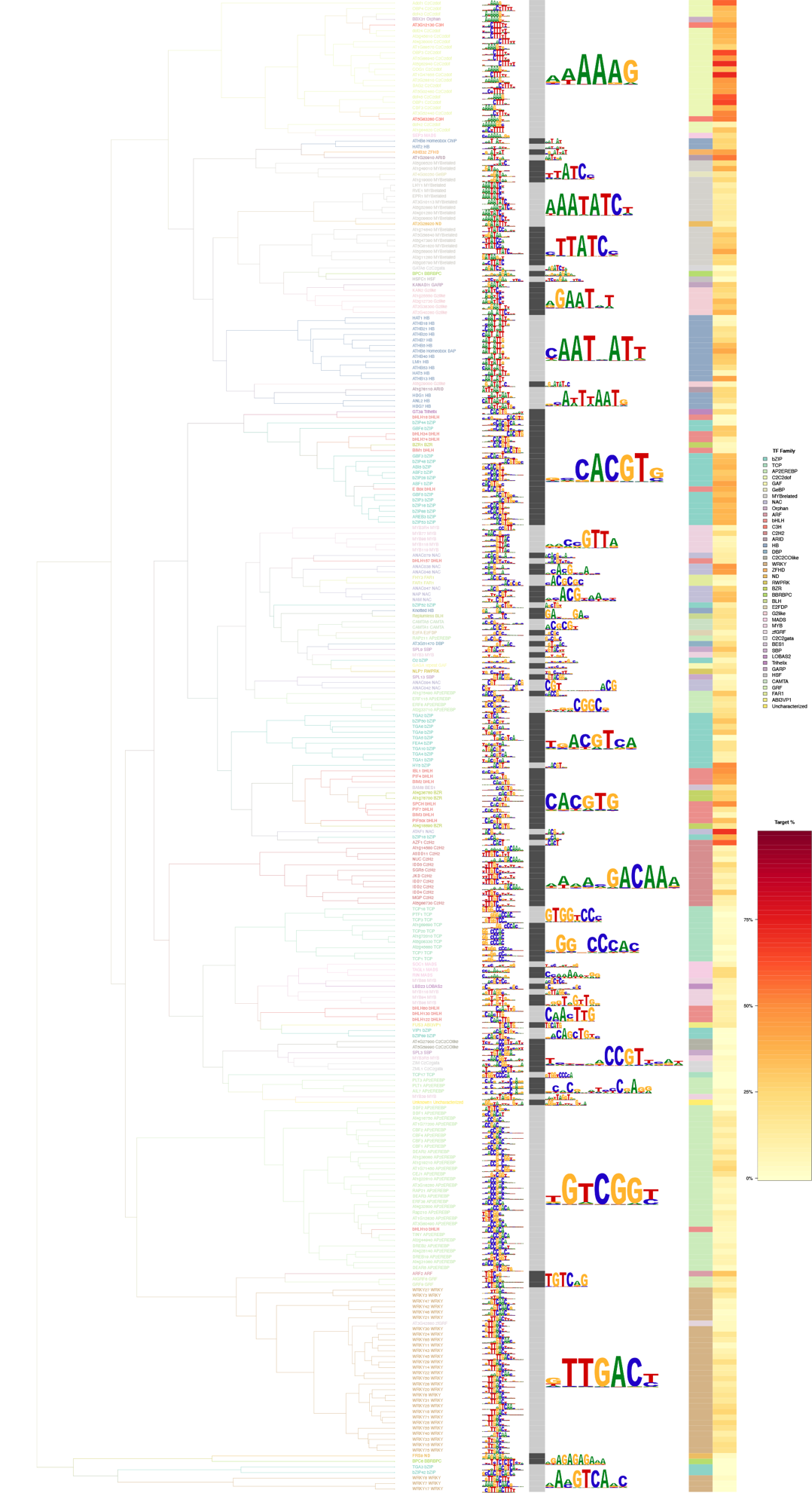

Figure S4. Motif tree on enriched motifs bound by ABI3 in *abi5* mutant background

Enriched motif tree found in DNA motifs from HOMER analysis in ChIP-seq of ABI3YHF/*abi5* mutants.

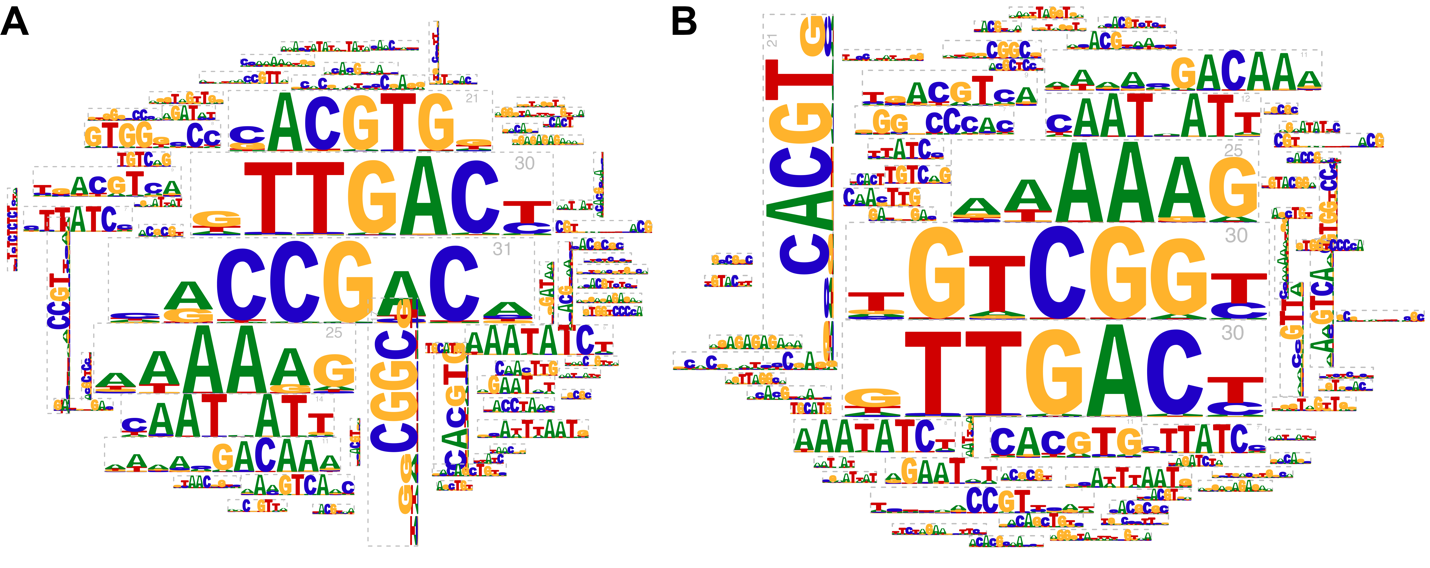

Figure S5. Motif cloud on enriched motifs bound by ABI3 in WT and *abi5* mutant backgrounds

Motif clouds found in DNA motifs from HOMER analysis in ChIP-seq of (A) ABI3YHF/WT and (B) ABI3YHF/*abi5* mutants.

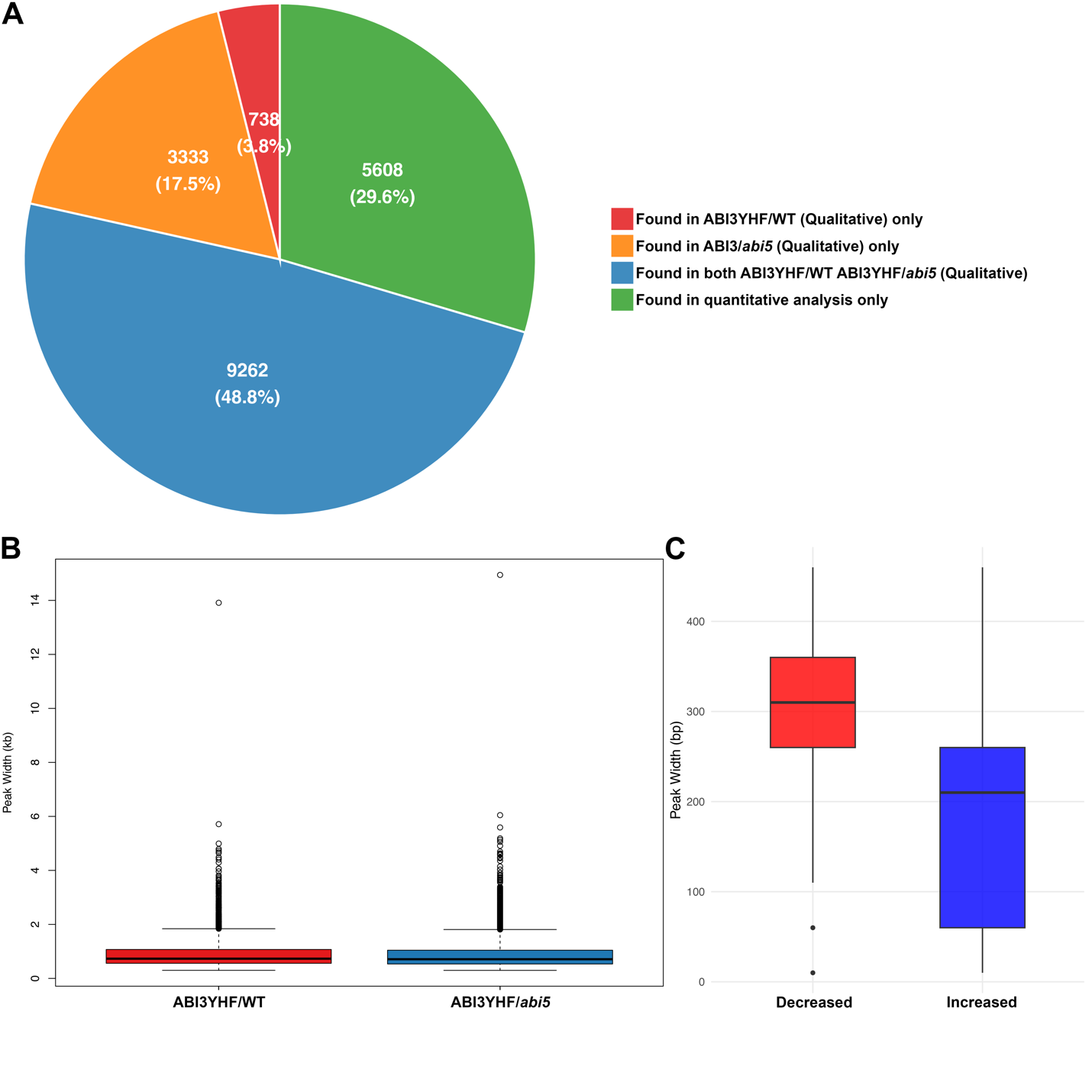

Figure S6. Quality of peaks analysed from csaw pipeline

(A) Pie chart on peaks annotated from csaw pipeline and MACS2-IDR pipeline. (B-C) peak width from (B) MACS2-IDR pipeline and (C) csaw pipeline in the form of box plot. Red color represents peaks from ABI3YHF/WT (B) and reduced binding in ABI3YHF/*abi5* (C) compared to ABI3YHF/WT. Blue color represents peaks from ABI3YHF/*abi5* (B) and increased binding in ABI3YHF/*abi5* (C) compared to ABI3YHF/WT.

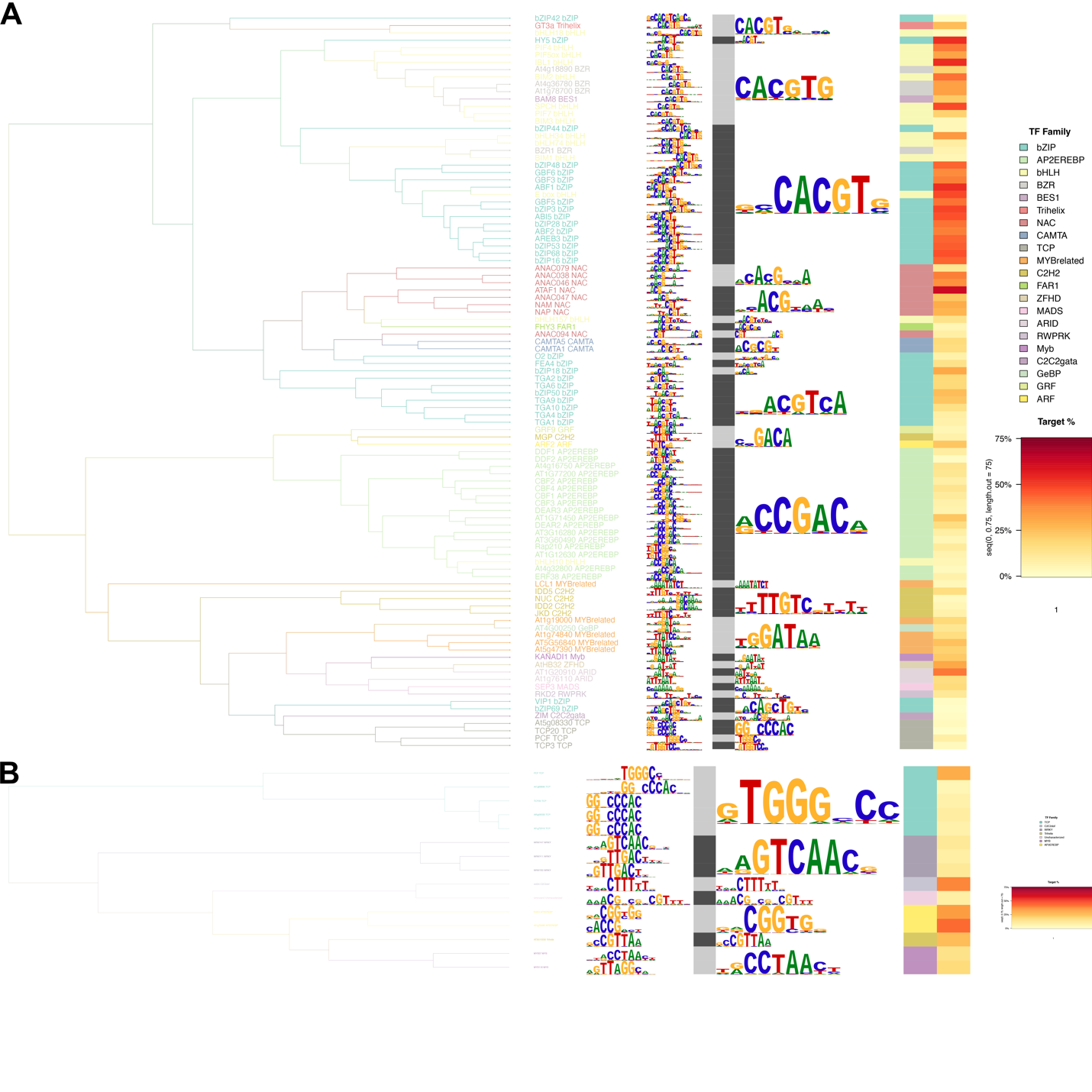

Figure S7. Motif tree on enriched motifs bound by ABI3 in abi5 mutant compared to WT background

Enriched motif tree found in DNA motifs from HOMER analysis in ChIP-seq of ABI3YHF/*abi5* mutants compared with ABI3YHF/WT with (A) decreased binding and (B) increased binding

**Supplementary Tables**

Table S1. PCR primers used for generating abi5 mutants

| Primer Name | Primer Sequence (5'-3') |
| --- | --- |
| ABI5_sg1_L_F | ATATATGGTCTCGATTGCGCAAGCGAGACATAATGGGTTTTAGAGCTAGAAATAGC |
| ABI5_sg2_L_R | ATTATTGGTCTCGAAACCTTTGTAGGAAGACTGTTGAAATCTCTTAGTCGACTCTAC |
| ABI5_sg3_L_F | ATATATGGTCTCGATTGAGTCTAGTCTTCCTCGACAGTTTTAGAGCTAGAAATAGC |
| ABI5_sg4_L_R | ATTATTGGTCTCGAAACCCGTTCTGAGCATTGTTCTGAATCTCTTAGTCGACTCTAC |

Table S2. PCR primers used for qPCR

| Primer Name | Primer Sequence (5'-3') |
| --- | --- |
| AT1G49240.1_ACT8_q1F | GTGTCTGGATTGGTGGTTCTAT |
| AT1G49240.1_ACT8_q1R | CTGGACCTGCTTCATCATACTC |
| AT3G24650.1_ABI3_q3F | CTGTGTCGGCTGAGGATTT |
| AT3G24650.1_ABI3_q3R | GCTGCTTCATCGCTTCTTTAC |
| AT2G36270.1_ABI5_q1F | CCGGTGTCTTCAGATGGATTAG |
| AT2G36270.1_ABI5_q1R | TCCTCTGTCTTCTCTCCACTAC |
| AT3G19920.2 _AtLDPS_q2F | AACTGGCTCCCACTGTTTAG |
| AT3G19920.2 _AtLDPS_q2R | AGTCGAGATGGCTTTGTCTATG |
| AT1G05510.1_AtOBAP1A_q3F | AGTTGCGAGAGGTTGACATAA |
| AT1G05510.1_AtOBAP1A_q3R | CATGATTCGAGGAGGGCTATAC |
| AT5G45690.1_AtOBAP2A_q1F | CGTGACGGATCAGAGAACTTAC |
| AT5G45690.1_AtOBAP2A_q1R | ATCATCGCCGCTCCTTTATC |
| AT1G29680.1_AtOBAP2C_q2F | ACTACTGGAAGCAACATGGTAAG |
| AT1G29680.1_AtOBAP2C_q2R | CATACAAGTGAGCCATGAGAGG |
| At5G16460.1_AtSEIPIN1_q1F | CGAGCTGACTGGTTCATGGT |
| At5G16460.1_AtSEIPIN1_q1R | CGGCGGTAAGAGCGGTATAG |
| AT4G25140.1_AtOLEO1_q1F | GGATTTACAAGTACGCAACG |
| AT4G25140.1_AtOLEO1_q1R | GTACGGTCACGGTCATGTTC |
| AT5G40420.1_AtOLEO2_q1F | GTACCCAAGTATTGTCCCTG |
| AT5G40420.1_AtOLEO2_q1R | CAAGCCCGATAGTTAGAGCC |
| AT3G01570.1_AtOLEO5_q1F | GCGTGTGTTTTGAGTGAACC |
| AT3G01570.1_AtOLEO5_q1R | TGTCTCCGACATTATTGTTACAGG |

Table S3 Binding preferences of ABI3YHF/WT and ABI3YHF/abi5 mutants

| Annotation | ABI3YHF/WT | ABI3YHF/abi5 |
| --- | --- | --- |
| Promoter (≤1 kb) | 77.10% | 74.90% |
| Promoter (1–2 kb) | 7.30% | 8.20% |
| Promoter (2–3 kb) | 2.70% | 3.20% |
| 5' UTR | 0.02% | 0.04% |
| Exon | 1.20% | 1.20% |
| Intron | 0.40% | 0.60% |
| 3' UTR | 4.00% | 3.70% |
| Downstream (≤300 bp) | 0.80% | 1.00% |
| Distal Intergenic | 5.60% | 6.30% |

Table S4 List of ABI5-independent ABI3 induced and repressed genes

| RNASEQ_RNA_UP_ABI5_UP | RNASEQ_RNA_DOWN_ABI5_UP |
| --- | --- |
| Locus | Locus |
| AT5G58575 | AT4G25730 |
| AT3G24515 | AT4G03430 |
| AT5G66100 | AT3G57000 |
| AT1G79010 | AT3G55200 |
| AT1G79340 | AT3G16770 |
| AT5G54780 | AT4G10070 |
| AT4G24940 | AT3G13380 |
| AT3G53740 | AT1G62120 |
| AT2G43360 | AT2G16090 |
| AT5G42750 | AT1G08115 |
| AT1G78180 | AT1G80150 |
| AT1G05562 | AT1G06070 |
| AT1G78730 | AT1G23980 |
| AT5G10620 |  |
| AT2G44650 |  |
| AT4G11130 |  |
| AT3G16950 |  |
| AT4G22830 |  |
| AT5G04660 |  |
| AT2G44210 |  |
| AT1G49500 |  |
| AT3G04910 |  |
| AT3G01900 |  |
| AT1G43780 |  |
| AT1G32120 |  |
| AT1G03780 |  |
| AT1G22070 |  |

Table S5 lipid body related genes bound by ABI3

| Symbol | geneId | Expression | foldChange | logFC | PValue | FDR | ABI5 Independent? |
| --- | --- | --- | --- | --- | --- | --- | --- |
| DGAT1 | AT2G19450 | 12361 | 1.28073613 | 0.35697327 | 1 | 1 | Independent |
|  |  |  | 1.3077888 | 0.38712958 | 1 | 1 | Independent |
| ASAT1 | AT3G51970 | 24 | 0.52032478 | -0.9425157 | 0.01433109 | 0.18132611 | Independent |
| PSAT1 | AT1G04010 | 126 | 0.68652837 | -0.5426088 | 1 | 1 | Independent |
| **SEIPIN1** | **AT5G16460** | **32** | **0.31163621** | **-1.6820652** | **6.68E-04** | **0.0168218** | **Dependent** |
| LDAP1 | AT1G67360 | 3598 | 1.631541 | 0.70623524 | 0.8182508 | 1 | Independent |
| LDAP2 | AT2G47780 | 7633 | 0.45626335 | -1.1320613 | 0.08694159 | 0.64139995 | Independent |
| LDAP3 | AT3G05500 | 2413 | 0.48521788 | -1.0432954 | 0.0218727 | 0.24616651 | Independent |
| LDIP | AT5G16550 | 404 | 1.2485235 | 0.32022297 | 1 | 1 | Independent |
|  |  |  | 0.664546 | -0.589559 | 0.5616242 | 1 | Independent |
| **OBAP1A** | **AT1G05510** | **4615** | **0.33421844** | **-1.5811368** | **2.04E-04** | **0.00656665** | **Dependent** |
|  |  |  | **0.41212721** | **-1.2788384** | **0.00296216** | **0.0532823** | **Dependent** |
| OBAP1B | AT2G31985 | 36 | 0.70007892 | -0.5144105 | 0.63665877 | 1 | Independent |
| **OBAP2A** | **AT5G45690** | **5060** | **0.19140843** | **-2.3852737** | **3.63E-07** | **4.07E-05** | **Dependent** |
|  |  |  | **0.18857293** | **-2.4068055** | **3.92E-08** | **6.63E-06** | **Dependent** |
| **OBAP2B** | **AT4G18920** | **136** | 0.40752193 | -1.2950504 | 0.03799633 | 0.36550966 | Independent |
|  |  |  | **0.38049762** | **-1.3940407** | **0.0031365** | **0.05578251** | **Dependent** |
| **OBAP2C** | **AT1G29680** | **463** | **0.29217699** | **-1.7750856** | **1.09E-05** | **6.21E-04** | **Dependent** |
| **OLE1** | **AT4G25140** | **1392** | **0.33922055** | **-1.5597045** | **0.00108288** | **0.02442** | **Dependent** |
|  |  |  | 0.44080535 | -1.1817864 | 0.0177403 | 0.21348136 | Independent |
| **OLE2** | **AT5G40420** | **673** | **0.14796677** | **-2.7566549** | **3.79E-10** | **1.12E-07** | **Dependent** |
|  |  |  | **0.16016464** | **-2.6423724** | **6.38E-10** | **1.73E-07** | **Dependent** |
| OLE3 | AT5G51210 | 226 | 0.84083564 | -0.2501043 | 1 | 1 | Independent |
|  |  |  | 0.78073101 | -0.3571025 | 1 | 1 | Independent |
| **OLE4** | **AT3G27660** | **169** | 1.69554117 | 0.76174581 | 0.36893923 | 1 | Independent |
|  |  |  | **0.28271548** | **-1.8225772** | **5.08E-06** | **3.48E-04** | **Dependent** |
| **OLE5** | **AT3G01570** | **1029** | **0.19856944** | **-2.3322845** | **1.69E-06** | **1.43E-04** | **Dependent** |
| OLE6 | AT1G48990 | 609 | 0.71774583 | -0.4784551 | 1 | 1 | Independent |
| OLE7 | AT2G25890 | 735 | 0.41384109 | -1.2728512 | 0.05517671 | 0.47547863 | Independent |
| **OLE8** | **AT3G18570** | **44** | **0.18115053** | **-2.464739** | **2.57E-09** | **6.16E-07** | **Dependent** |
| CLO1 | AT4G26740 | 176 | 0.48303226 | -1.0498086 | 0.04632081 | 0.42223036 | Independent |
| **CLO2** | **AT5G55240** | **90** | **0.17556668** | **-2.509909** | **5.97E-10** | **1.70E-07** | **Dependent** |
| HSD2 | AT3G47350 | 1734 | 1.11887173 | 0.16204465 | 0.70075096 | 1 | Independent |
| HSD5 | AT4G10020 | 579 | 0.53251392 | -0.9091089 | 0.24929704 | 1 | Independent |
|  |  |  | 0.54515959 | -0.8752495 | 0.15732909 | 0.89835419 | Independent |
| **HSD6** | **AT5G50770** | **50** | **0.37660447** | **-1.408878** | **0.00125611** | **0.02715983** | **Dependent** |
| CAS1 | AT2G07050 | 2494 | 0.85278977 | -0.229738 | 1 | 1 | Independent |
| LIDL1 | AT1G18460 | 758 | 0.41931611 | -1.2538898 | 0.08548536 | 0.63397738 | Independent |
|  |  |  | 0.43620435 | -1.196924 | 0.00918182 | 0.1309612 | Independent |
| LDAH1 | AT1G10740 | 1255 | 1.18792212 | 0.24844026 | 1 | 1 | Independent |
| **LDDH2** | **AT1G19400** | **406** | 0.65201966 | -0.6170126 | 0.13723139 | 0.83498225 | Independent |
|  |  |  | **0.28193433** | **-1.8265689** | **5.77E-06** | **3.80E-04** | **Dependent** |
| SLDP1 | AT1G65090 | 1611 | 0.59392172 | -0.7516553 | 0.42236247 | 1 | Independent |
|  |  | 1611 | 0.46999748 | -1.0892751 | 0.06418384 | 0.52489415 | Independent |
| **SLDP2** | **AT5G36100** | **511** | **0.34554522** | **-1.5330536** | **6.12E-04** | **0.01579888** | **Dependnent** |
|  |  |  | 0.68659488 | -0.542469 | 1 | 1 | Independent |
| LIPA | AT1G07985 | 3473 | 0.32794389 | -1.6084791 | 0.01524691 | 0.19074755 | Independent |
| **LDPS** | **AT3G19920** | **1755** | 0.44944152 | -1.1537947 | 0.02631378 | 0.28014407 | Independent |
|  |  |  | **0.35680135** | **-1.486807** | **2.86E-04** | **0.00853813** | **Dependnent** |
|  |  |  | 1.16653431 | 0.22222874 | 1 | 1 | Independent |
|  |  |  | 1.19785489 | 0.26045315 | 1 | 1 | Independent |
| ERD7 | AT2G17840 | 876 | 0.49200011 | -1.0232695 | 0.06217191 | 0.51302797 | Independent |
|  |  | 876 | 0.48595731 | -1.0410985 | 0.05616858 | 0.48032168 | Independent |
| SDP1 | AT5G04040 | 2924 | 0.80632652 | -0.3105639 | 1 | 1 | Independent |
|  |  |  | 1.17689131 | 0.23498109 | 1 | 1 | Independent |
| SDP1-LIKE | AT3G57140 | 1074 | 1.61379548 | 0.69045775 | 0.39331499 | 1 | Independent |
|  |  |  | 1.21864825 | 0.28528177 | 1 | 1 | Independent |
| OBL1 | AT3G14360 | 491 | 0.71569361 | -0.482586 | 1 | 1 | Independent |
|  |  |  | 0.68967095 | -0.5360199 | 1 | 1 | Independent |
